# Adaptive introgression during environmental change can weaken reproductive isolation

**DOI:** 10.1101/553230

**Authors:** Gregory L. Owens, Kieran Samuk

## Abstract

Anthropogenic climate change is an urgent threat to species diversity. One aspect of this threat is the collapse of species reproductive barriers through increased hybridization. The primary mechanism for this collapse is thought to be the weakening of ecologically-mediated reproductive barriers, as demonstrated in many cases of “reverse speciation”. Here, we expand on this idea and show that adaptive introgression between species adapting to a shared, moving climatic optimum can readily weaken *any* reproductive barrier, including those that are completely independent of climate. Using genetically explicit forward-time simulations, we show that genetic linkage between alleles conferring adaptation to a changing climate and alleles conferring reproductive isolation (intrinsic and/or non-climatic extrinsic) can lead to adaptive introgression facilitating the homogenization of reproductive isolation alleles. This effect causes the decay of species boundaries across a broad and biologically-realistic parameter space. We explore how the magnitude of this effect depends upon the rate of climate change, the genetic architecture of adaptation, the initial degree of reproductive isolation, the degree to which reproductive isolation is intrinsic vs. extrinsic, and the mutation rate. These results highlight a previously unexplored effect of rapid climate change on species diversity.

## Introduction

Global climate change (GCC) is expected to be an increasingly important stressor in the next century (Thomas et al. 2004, Hoffmann and Sgro 2011). GCC can lead to significant fitness costs or extinction if populations cannot adjust their ranges (e.g. via migration) or adapt to the changing climate (Jump and Penuelas 2005; Zimova et al. 2016). Apart from its direct effects on species fitness, GCC can also profoundly alter the basic ecological functioning of ecosystems e.g. via changes in phenology or species composition (Walther et al. 2002). As such, GCC represents an existential threat to biological diversity, at the level of populations, species, and ecosystems. Considering the central role of biological diversity in the functioning of evolutionary and ecological processes, understanding the full biological effect of GCC remains a key problem.

One potential effect GCC is increased interspecies hybridization, which can itself precipitate further loss of biodiversity (Rhymer and Simberloff 1996, Todesco et al. 2016). Such hybridization can cause a common species to subsume a rare species (Oliveira et al. 2008; Beatty et al. 2014; Vallender et al. 2007) or the collapse of multiple species into a single hybrid swarm (Taylor et al. 2006). In both cases, genetic and species diversity are lost. GCC may also precipitate the collapse of species boundaries by breaking down spatial, temporal or behavioural premating barriers (Chunco 2014). At the simplest level, range shifts can bring together species that may have no other significant barriers (e.g. Garroway et al. 2010). Reproductive barriers relying on the timing of life-history events are also susceptible to climate change which can increase the temporal overlap between species (e.g. Gerard et al. 2006). Furthermore, behavioural isolation may rely on environmental cues disrupted by GCC, as in spadefoot toads in which water conditions influence mate choice (Pfennig 2007). The breakdown of these barriers may lead to loss of more rare species under GCC, or the collapse of sister species, as has been seen in smaller localized environmental shifts (Taylor et al. 2006; Vonlanthen et al. 2012).

While perhaps unfamiliar to many workers in the field of climate change science, there is a rich theoretical literature dedicated to the study of the dynamics of interspecific hybridization and gene flow (reviewed in e.g. Abbot 2013 and Seehausen 2013). Often framed in the context of the evolution and maintenance of reproductive isolation (i.e. speciation), this literature has provided many key insights germane to the study of climate change induced hybridization. For example, hybrid zone models have shown that universally adaptive alleles readily introgress across hybrid zones, while alleles that cause reproductive isolation generally resist introgression (Barton 1979, Gompert et al. 2012, Barton 2013). Other modelling efforts have revealed that depending on the balance between reproductive isolation and shared ecological selection, introgression can cause previously isolated populations to remain isolated or collapse into hybrid swarms (e.g. Buerkle 2000). Further, the balance between divergent natural selection, gene flow and reproductive isolation has been extensive explored in the theoretical speciation literature (see Barton 2013 and references therein). However there has thus far been poor integration between models of reproductive isolation and models of adaptation to climate change.

The fact that introgression can transfer alleles between species has led to the idea that hybridization could facilitate adaptation to GCC through the transfer of adaptive alleles between species, i.e. adaptive introgression. This has traditionally been studied in the context of species/populations with pre-existing differential adaptation to the changing climate variable; for example a warm adapted species transfering alleles to a cold adapted species (e.g. Gómez et al. 2015). In this example, one species acts as a pool of alleles preadapted to a future climatic optimum. Importantly, in these types of models, introgression is being driven by selection and not demographic processes or perturbations of prezygotic isolation, as seen in other models where climate change drives hybridization.

What has not been appreciated in previous models of adaptation to a changing climate is that during a rapid environmental shift, segregating variation within two reproductively isolated species could theoretically undergo adaptive introgression even if neither species is particularly preadapted to the environmental shift. We propose that climate-induced adaptive introgression could readily occur in most species because (1) the identity of the particular alleles involved in climatic adaptation are likely idiosyncratic in each species/population, and (2) these alleles could, in principle, be globally adaptive under a GCC scenario. Indeed, segregating climate adaptation alleles (or linked blocks of alleles) could easily be strong enough to outweigh the fitness costs of any linked reproductive isolation alleles. As a side effect, reproductive isolation alleles could readily be homogenized between species, reducing reproductive isolation and precipitating the collapse of species boundaries. This scenario dramatically increases the likelihood of GCC-induced introgression from populations differing in altitude or latitude, to nearly any parapatric pair capable of hybridization, even if reproductive isolation is initially high.

Here, we direct testly of the role of climate-induced adaptive introgression in degrading reproductive barriers using state-of-the-art forward time population genetic computer simulations. We envision a scenario in which two species are initially reproductively isolated (but capable of limited hybridization) and must cope with an extreme shift in the environment. We explore the parameter space under which adaptation to a shifting climatic optimum drives a reduction in reproductive isolation. We interpret these results in the context of the future anthropogenic climate change and the loss of species diversity.

## Methods

### Conceptual model

We consider the scenario of two parapatric species inhabiting demes in two different habitats. These species exchange migrants at a low level, but reproductive isolation via local adaptation (i.e. extrinsic postzygotic isolation and immigrant inviability) is strong enough to prevent substantial introgression. We imagine that these two species must also cope with constant adaptation to a shared oscillating “climate” optimum. This climatic optimum does not not directly affect the degree of local adaptation and/or reproductive isolation, i.e. reproductive isolation is completely independent of the direct effects of climate. The climate oscillation continues for a long initial burn in period, during which alleles conferring adaptation to climate accumulate in each species. After this period, the oscillation ends and the climatic optimum begins rapidly increasing at a constant rate, as is expected under projections of anthropogenic climate change.

We hypothesize that if the rate of change in the climatic optimum is sufficiently high, selection for migrant alleles conferring increased climate tolerance will overwhelm the negative fitness effects of linked reproductive isolation alleles. This will cause the erosion of reproductive isolation between species and increase the chance of speciation reversal. Importantly, we expect this outcome even when the strength of ecological selection mediating reproductive isolation itself is orthogonal to the strength of climate-mediated selection.

### Model details

We implemented the above conceptual model as a genetically explicit Wright-Fisher model in SLiM 3.0 (Haller and Messer 2018). As in all Wright-Fisher models, population sizes are constant, all fitness is relative and extinction is impossible. The details of our implementation are depicted graphically in Figure 1 and a list of simulation parameters and their values are detailed in Table 1. We simulated two diploid populations of constant size *Ne*, with a constant migration rate of *m* proportion migrants per generation. Each individual was initialized with 99999 genetic loci contained on a single chromosome with a uniform recombination rate of *r* between loci. We initially scaled the recombination rate so that the entire genome was 100 cM in length, but also explored varying recombination rates up a genome size of 1000 cM (see Results). We modelled local adaptation in the two populations as *l*_*EX*_ divergently selected alleles at loci evenly spaced across the chromosome, with each population fixed for a different allele. Divergently selected alleles imposed a fitness cost of *s*_*RI*_ when not found in their home population/habitat, modelling extrinsic postzygotic isolation.

**Table 1.**
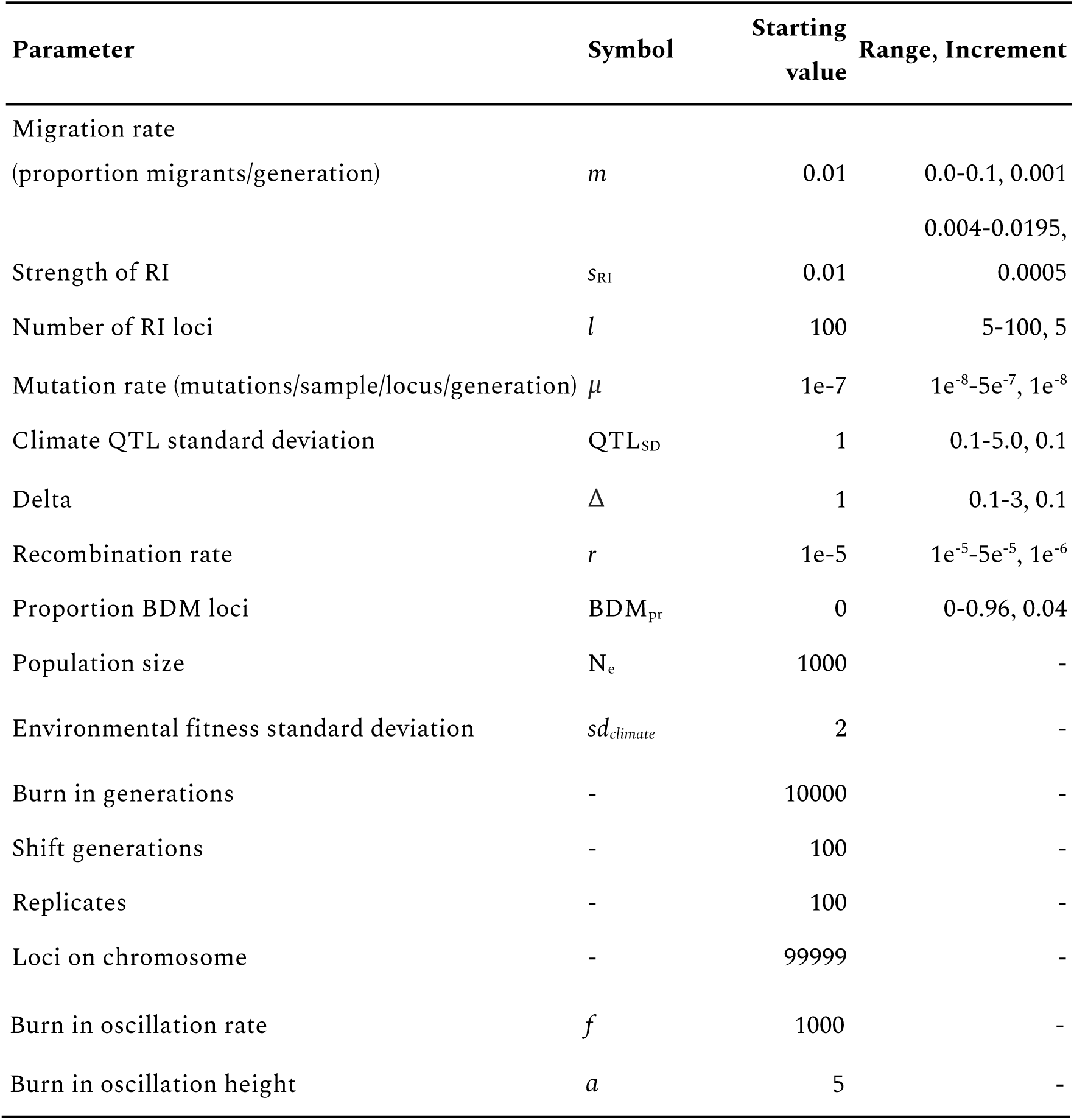
Parameters of the adaptive introgression simulations. For each set of simulations, each parameter was set to the starting value which was varied from the minimum to maximum value by the specified increment.

**Figure 1.**
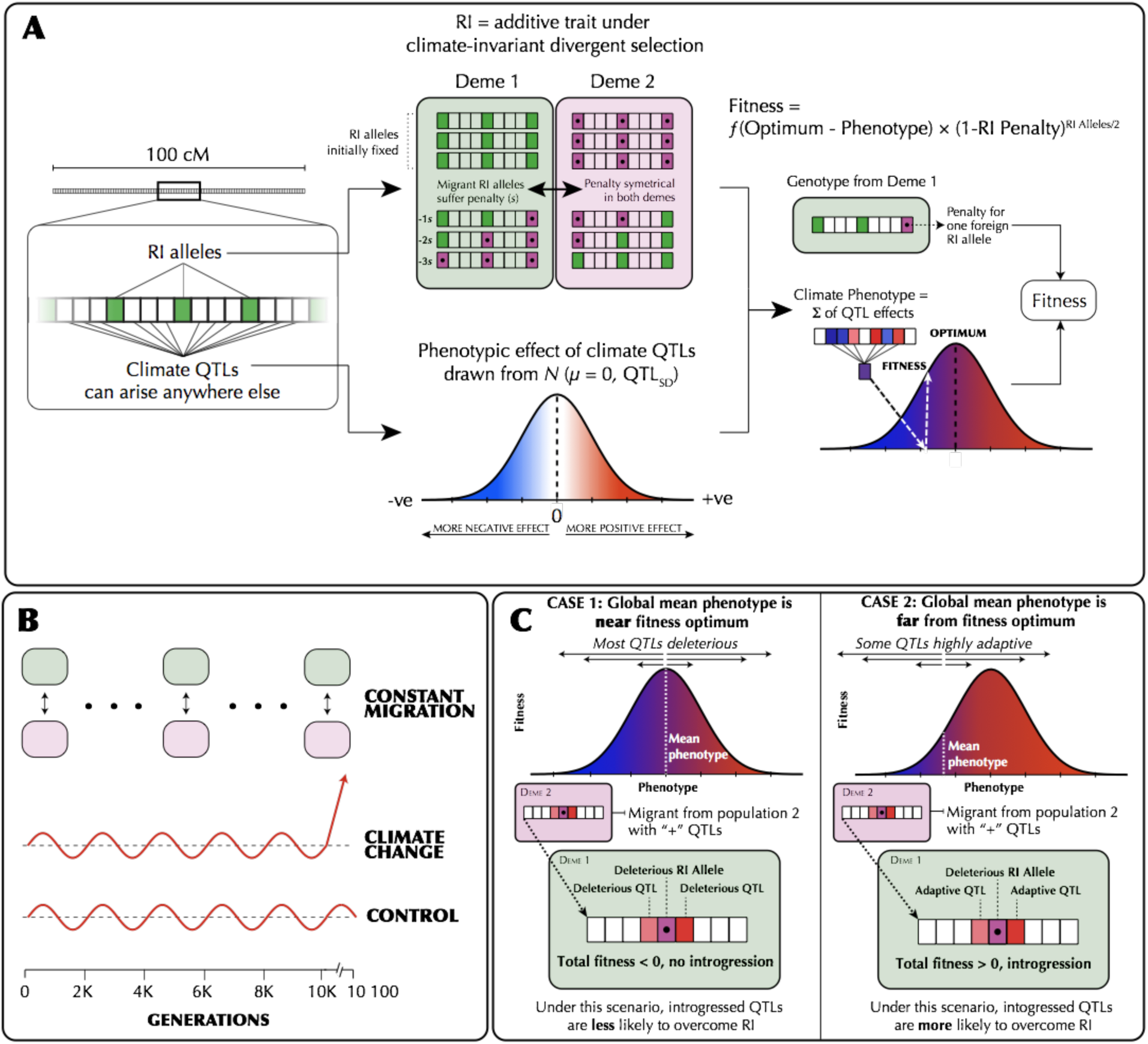
(A) The genetic architecture of adaptation and speciation in the model. From left to right: Each individual has a single 100-1000 cM chromosome, over which reproductive isolation (RI) loci occur at regularly-spaced intervals. These loci are initialized with RI alleles (at 100% frequency) that confer local adaptation to one of two initial demes (depicted as green or purple/dotted alleles, corresponding to Deme 1 or Deme 2 environments). Both demes are of equal size (*Ne* = 1000). All non-RI loci (depicted as white/transparent loci, initially) have the potential to give rise to climate-adaptation alleles. The phenotypic effects of each these alleles are drawn from a normal distribution (shown as a gradient from blue to white to red). An individual’s climate phenotype is the sum of the phenotypic effects of its climate QTLs (pure additivity). The fitness of each individual is a function of the number of foreign RI alleles and the phenotypic distance of that individual from the environmental optimum, with the climate fitness landscape modelled as a gaussian distribution (shown as a gradient from blue to red). (B) The course of the simulation. Migration rate and population size of the two demes is held constant. In each replicate simulation, the fitness optimum fluctuates regularly for a 10 000-generation burn-in period. The state of the initial population is then duplicated and subjected to 100 additional generations of (1) a climate change scenario in which the climatic optimum rapidly shifts in a single direction and (2) a control scenario in which the optimum continues its fluctuation course. (C) The conditions under which adaptive introgression overwhelms RI. On the left, if the two populations are able to individually track the climatic optimum, newly-arising climate alleles are only able to exert either weakly positive or (more commonly) negative effects on fitness due to overshooting the optimum. In contrast, on the right, if the populations cannot effectively track the optimum, there is scope for climatic alleles to have large positive fitness effects. If these fitness effects are sufficiently large, these alleles can overwhelm the negative fitness effects of linked RI alleles and introgress between demes, degrading overall reproductive isolation.

In addition to extrinsic postzygostic isolation, we also modelled intrinsic postzygotic isolation using two-locus Bateson-Dobzhansky-Muller incompatibilities (Bateson 1909, Dobzhansky 1936, Muller 1942). These epistatic incompatibilities were modelled as a fitness cost of *s*_*RI*_ scaled by the number of negatively-interacting pairs of alleles from each population (see Supplementary Appendix 1 for details). When testing the effects of BDMs, we maintained a constant number of total reproductive isolation loci, but varied the proportion of loci that were extrinsic or BDM loci (*l)*. We also explored the effect of the total number of RI loci (i.e. the genetic architecture of RI *per se*) on the potential for adaptive introgression/hybridization. To keep the total magnitude of RI similar between simulations, we always co-varied *s*_*RI*_ so that the *s*_*RI*_ x *l* was held constant. To allow for fine-scale view of introgression, we tracked ancestry was using 100 neutral alleles initially fixed between the populations, spread evenly across the genome. All alleles of selective/phenotypic effect were codominant with dominance = 0.5.

In addition to reproductive isolation, individual fitness also depended on their phenotypic distance from a climatic optimum. This optimum was initially 0, and during the burn in period oscillated from −5 to 5 (in arbitrary units) every 500 generations based on the formula: *sin*(π \**generation* / 500) /5. The individual phenotype was determined by alleles at QTL-like climate loci which could appear via mutation at all sites other than RI or ancestry tracking loci (i.e. 99899 - *l* sites). Climate QTL mutations occurred at a rate *μ* per locus per sample per generation and their phenotypic effect was drawn from a gaussian distribution with a mean of zero and a standard deviation of *QTL*_*SD*_. Conceptually, these QTL climate alleles modify whether an individual is “hot” (positive effects) or “cold” (negative effects) adapted.

The first step of the simulations was a burn-in of 10*Ne* generations to simulate the generation of standing genetic variation under normal climatic conditions. At the end of the burn-in period, the complete state of each replicate simulation was saved. Each simulation was then continued under both a “control” and climate change scenario for an additional 100 generations. In the control scenario, the environmental oscillation continued as normal. In contrast, under the climate change scenario the phenotypic optimum increased by a rate of Δ each generation without oscillation. In each generation we recorded the average degree of reproductive isolation, mean fitness, the mean and standard deviation of the climate phenotype and the amount of introgressed ancestry for each population. Reproductive isolation (RI) was calculated accounting for the extrinsic and BDM loci. For extrinsic loci, RI was the difference in fitness for an average individual in their home habitat vs. the foreign habitat. For BDMs, since fitness penalties occur only in F1 hybrids and beyond, we calculated the expected average magnitude of BDM fitness costs based on Hardy-Weinberg expectations in F1s (Supplementary Appendix 1). Finally, for each simulation we report the mean introgressed ancestry and reproductive isolation between the start and end of control and test scenarios, as well as the mean rate of phenotypic change in Haldanes for the test scenario. A Haldane is a measure of evolutionary change in log mean trait value in units of standard deviation of that log trait (Gingerich 1993). All formulas used in the simulation are presented in the Supplementary Appendix and all code for underlying simulations is available at https://github.com/owensgl/adaptive_introgression.

To explore the parameter space under which adaptive introgression mediates RI collapse, we systematically varied the following parameters: mutation rate (*μ*), migration rate (*m*), strength of divergent selection (*s*_*RI*_), the number of divergently selected loci (*n*_*RI*_), the proportion of BDMs (*pr*_*BDM*_),the standard deviation of QTL effect sizes (*QTL*_*SD*_) the recombination rate (*r*), and the rate of climate change (Δ). We varied each parameter independently and kept the other parameters at a default value known to permit a low level of introgression in preliminary tests (Table 1). Each parameter set was replicated 100 times. All analyses were carried out in R 3.5.1 (R Core Team 2018) and plotting was done using ggplot2 (Wickham 2016).

Finally, while our primary goal was testing the detrimental effects of hybridization, we also examined the potential *beneficial* effects of climate change induced introgression, i.e. to what degree introgression facilitates adaptation. To do this, we ran simulations varying the rate of climate change with (*m*=0.01) or without (*m*=0) migration. At the last generation (gen=10,100), we compared the average climate phenotype to the current phenotypic optimum. We defined “adaptational lag” as the difference between these values divided by the rate of climate shift. This represents how many generations behind the current generation that the population is adapted to. For example, assume the optimum increases by 2 per generation and is currently 100, if the average phenotype is 90, then the adaptational lag is 5 (e.g. (100 - 90) / 2).

## Results

### Rapid climate change and adaptive introgression facilitates collapse of species boundaries

When climate change is rapid, we find that adaptive introgression of climate QTL alleles rapidly drives the homogenization of allele frequencies at linked RI loci between species. Figure 2 visualizes one example simulation where after 100 generations of climate change, RI is degraded to nearly half its original strength (Figure 2A) and introgressed climate QTL alleles are common (Figure 2B). As climate QTL alleles move between populations, RI and neutral alleles hitchhike along with them resulting in substantial genome-wide introgression (Figure 2A & 2C). In contrast, in the control scenario without climate change, RI remains intact and introgression is minimal (Figure 2 E-H).

**Figure 2.**
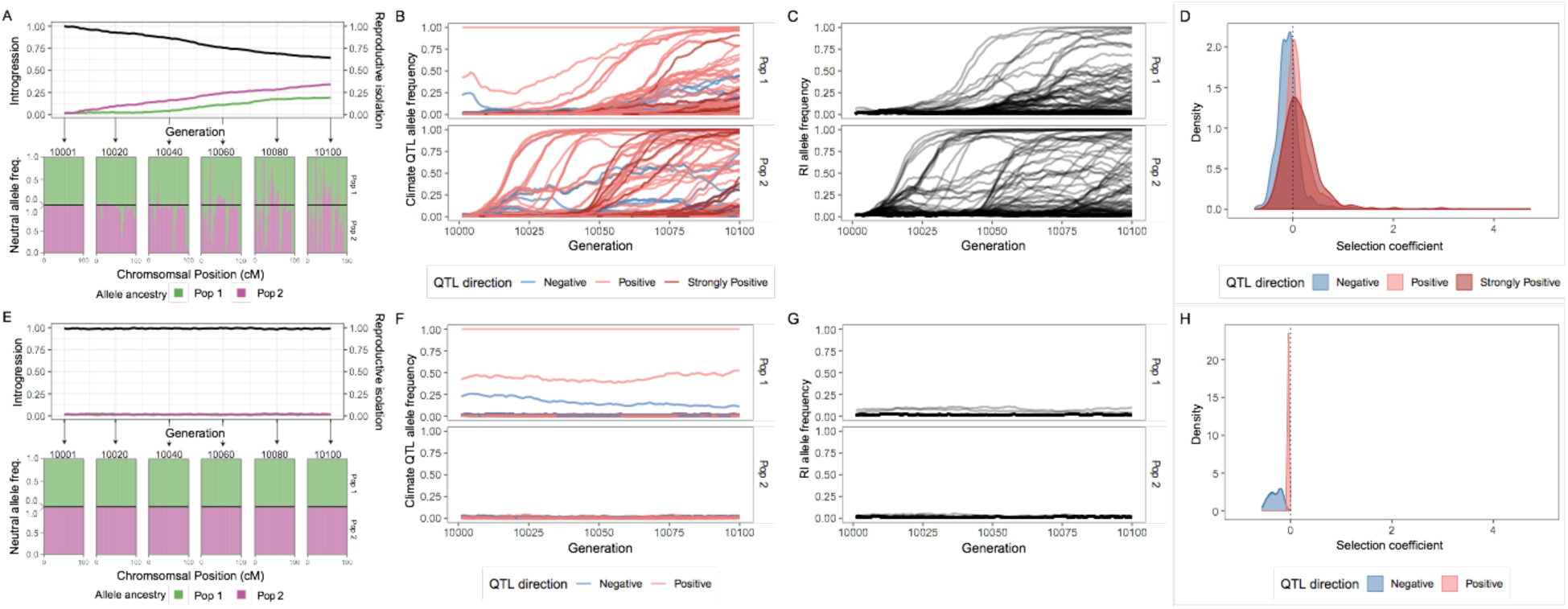
A single example simulation with Δ = 1.5, illustrating the climate driven adaptive introgression. Panels A-D present the test climate change scenario, while E-H are the control scenario. (A & E) The upper half is the average introgressed ancestry for each population (purple and green) and the average reproductive isolation between populations (black). The lower half is the ancestry for neutral loci during the post-burn in period at 20 generation intervals. The top and bottom parts of this portion represent population 1 and 2 respectively. (B & F) The allele frequency trajectory for introgressed climate QTL color coded by QTL strength. Color codes QTL effect; -ve phenotypic effect (blue), +ve effects (light red) or large +ve effect (>2, dark red). (C & G) The allele frequency trajectory for introgressed RI alleles (D & H) The distribution of selection coefficients on QTL loci per population per generation. Color groups represent QTL with -ve phenotypic effect (blue), +ve effects (light red) or large +ve effect (>2, dark red). Plot is filtered to only include loci with allele frequency < 0.9 and > 0.1.

For a wide range of parameters we find decreased reproductive isolation and increased introgressed ancestry under the climate change scenario (Figures 3 and S1). This effect is enhanced by reduced levels of genetic variation; both reducing mutation rate and reducing the standard deviation of the climate QTL effect size increases the likelihood and magnitude of adaptive introgression (Figure 3A, Figure 3D). Increased migration generally causes increased RI loss, although high migration degrades RI even in the absence of climate shifts (Figure 3C). Interestingly, increasing the average fitness effect of RI loci has minimal effect on the amount of RI lost, although below a threshold, populations merge and all RI is lost during the burn in period (Figure 3G). Recombination rate has two effects; during the burn in, increased recombination allows for more RI loss, while during climate change decreased recombination leads to slightly more RI loss (Figure 3F). When varying the genetic architecture of RI, we find that fewer strong RI loci, lead to less RI loss (Figure 3H). We find that intrinsic isolation is much less effective at maintaining RI during the burn in period, consistent with previous simulations (Bank et al. 2012). Climate change increases the amount of introgression and RI loss when intrinsic isolation is present unless populations are already completely merged (Figures 3E, S1E). Lastly, increasing the rate of climate change in the test scenario increases the amount of adaptive introgression (Figure 3B).

**Figure 3.**
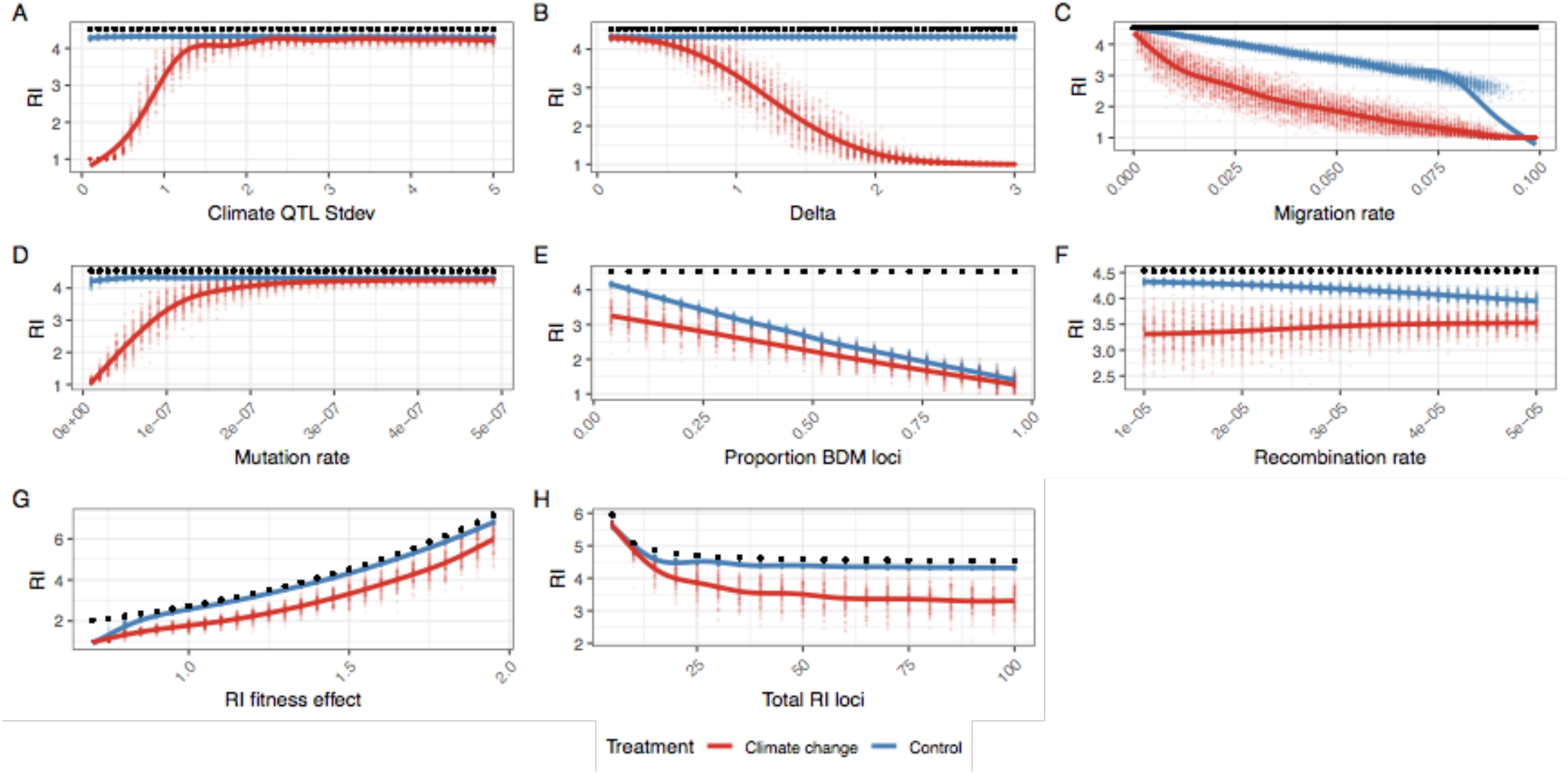
The average reproductive isolation at generation 10,100 for climate change (red) and control simulations (blue), while varying individual parameters. RI is defined as the home fitness advantage which is the fold fitness advantage for the average sample in its home environment compared to the alternate environment based on divergent selection and BDM loci. A value of 1 means equal fitness in both environments and there is no RI. The black dot is the initial and maximum level of RI for each simulation. Individual parameters were varied to show the effect of (A) climate QTL effect size standard deviation, (B) optimum shift per generation (delta), (C) migration rate, (D) climate QTL mutation rate, (E) proportion of RI loci that are BDM instead of extrinsic, (F) the recombination rate, (G) the fitness effect of each RI loci and (H) the number of RI loci.

### Simulation parameters fall within realistic ranges

To better connect our simulation results with observations from natural populations, we measured the rate of phenotypic evolution in the test scenario in Haldanes (standard deviations per generation). We find that most simulations had average evolutionary rates ranging from 0.01 to 0.06 Haldanes and are well within the range of empirical estimates from natural populations (Hendry et al. 2008). That said, in simulations with very high rates of climate change (Δ>2.5), we found large and erratic evolutionary rates. For many of these simulations, one or both species failed to track the changing environment causing fitness of all individuals to fall to zero and end the simulation.

In example simulations, we found that 99.3% of the realized selection coefficients for introgressed climate QTL alleles fell below 1 (Figure 2D). QTL alleles with positive and strongly positive phenotypic effects were more likely to have positive selection coefficients, consistent with the importance of the climate optimum.

### Beneficial effects of introgression

We find that migration can reduce adaptational lag when climate change is moderate or high (Figure 4). This effect is particularly strong when climate change is rapid.

**Figure 4.**
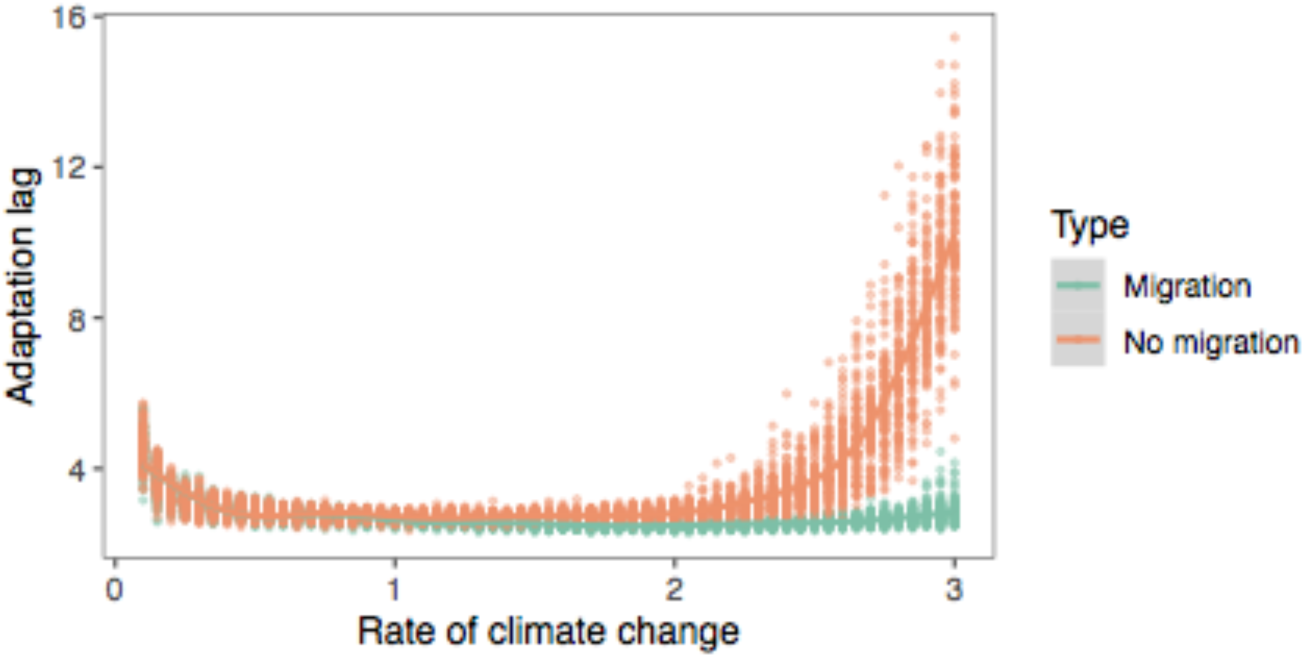
The adaptational lag at the final generation for simulations with (m=0.01) and without migration (m=0). Adaptational lag is defined as the phenotypic optimum minus the phenotypic mean divided by the rate of change in optimum, and represents how many generations behind the changing optimum that the population is.

## Discussion

Global climate change is expected to have wide-ranging detrimental effects on biodiversity. The movement of beneficial alleles between species, i.e. interspecific adaptive introgression, is likely to increasingly relevant as the climate warms. In spite of increased interest in adaptive introgression per se, we still have a poor understanding of the incidental effects of adaptive introgression on the integrity of species boundaries. Our simulations have shown that it is possible for adaptation to a common, changing environment to cause introgression and speciation reversal. Importantly, we observed this effect in scenarios where the mechanism of RI is itself completely independent of the changing climate.

### Will adaptive introgression lead to speciation reversal?

In our simulations, several parameters had a strong effect on the amount of introgression. When adaptive variation is limited, RI is initially weak, or environmental change is rapid, complete genetic homogenization is likely. In these cases, RI is completely degraded and would clearly represent speciation reversal in a natural system. In more moderate parts of parameter space, introgression is increased during environmental change, but populations do not completely homogenize (Figure S1). In these cases, RI is still eroded between populations (Figure 3). Importantly, we believe our estimates of RI loss are likely somewhat conservative, because we do not include any additional factors that would contribute to species collapse (e.g. cases where RI is directly affected by a change in climate).

We found that in the absence of divergent selection intrinsic epistatic isolation (BDM incompatibilities) were unable to maintain RI during the burn in period. This result is consistent with previous modelling of parapatric speciation (Barton & Bengtsson 1986, Bank et al. 2012, Lindtke & Buerkle 2015). Consistent with their effect in the burn in period, during climate change, introgression and RI loss is enhanced when intrinsic RI is present. Thus, although we have focused on extrinsic RI, intrinsic RI is also susceptible to adaptive introgression.

The ultimate question of which species are in danger of reverse speciation is dependent on a multitude of interacting factors and is beyond the scope of this paper, but we can highlight several risk factors.

1. For hybridization to be an issue, a potential hybridizing species must be at least in parapatry. Surveys have estimated the percent of species that hybridize with at least one other congener to be around 10-25%, although if climate change disrupts species ranges or premating isolation, that number may increase (Mallet 2005).
2. The rate of environment change and the steepness of the changing fitness landscape. Species with broader climate niches will be less susceptible because they will be under weaker selection.
3. The genetic architecture of climate adaptation in the species. Species with numerous large effect climate adaptation alleles segregating within their gene pool will be more able to adapt to the changing climate *without* introgressed alleles. Low diversity species will be more susceptible and reliant on adaptive introgression.
4. The genetic architecture of reproductive isolation between species. Species with few large effect RI loci will be more resistant to RI decay than species with a more diffuse and polygenic RI architecture.
5. The demographic and life history of the species. Unbalanced population sizes may result in one population harboring more adaptive alleles and lead to unbalanced introgression. Small populations will also be more susceptible to extinction due to the fitness costs of introgressed RI alleles. Features that reduce effective population size, e.g. high variance in reproductive success, are also likely to have reduced diversity of climate adapting alleles.

### Implications for the fate of species in changing environments

Our simulations suggest that rapidly changing environments can cause the collapse of species barriers in the *absence* of any direct effect on the underlying strength of reproductive isolation. By design, we modelled a scenario in which the strength of RI (modelled as divergent selection) is (a) invariant throughout (i.e. not reduced by environmental change itself) and (b) totally orthogonal to the strength of climate-mediated selection (i.e. extrinsic RI alleles do not affect the climate phenotype). This is an important departure from previous work, in which the collapse of reproductive isolation or “reverse speciation” occurs because RI is itself dependent on the environment (e.g. trophic or sensory niche).

This difference has several important implications. For one, the mechanism we outline here can occur in any population where adaptive introgression is possible (i.e. RI is not absolute and the climate-mediated selective optimum is to some degree shared). This greatly expands both the number of populations that may be susceptible to introgressive collapse and the potential severity of such collapses. For example, adaptive introgression could act in concert with the collapse of climate-mediated reproductive barriers, accelerating collapse. Further, while we focused on extrinsic isolation, our simulations show that scenarios where reproductive barriers are independent of ecological context (e.g. in the case of purely intrinsic isolation) are not immune.

Although we have framed our discussion in the context of climate change, our results are applicable to any strong, consistent, and shared selective event. These events include any environmental or ecological disturbance that alters the shared selective landscape of the two populations such that both populations are sufficiently displaced from their selective optima (thereby increasing the average size of selection differentials between genotypes, i.e. the strength of selection). One such event that has been studied in natural systems is eutrophication, which has been suggested to have caused speciation reversal in European lake whitefish (Vonlanthen et al. 2012). Thus far, this reversal has been attributed to changes in RI as a direct result of ecological and/or behavioural changes. However, if eutrophication exerts a common selective pressure on a group of parapatric species (e.g. mediated through changes in water chemistry) introgression could become adaptive and contribute to the collapse of species boundaries. Similarly, ocean acidification could be a strong source of shared selection and may induce introgression between previously well isolated species (Pespeni et al. 2013).

In contrast to our hypothesis of introgression driving species collapse, adaptive introgression of an insecticide-resistance mutation in *Anopheles* mosquitoes lead to the homogenization of previously differentiated genomic region, but not genome wide despeciation (Clarkson et al. 2014). In this example, selection seemingly only acts on a single large effect locus, rather than the polygenic architecture of climate adaptation we hypothesize. This limits the amount of introgression in comparison to our simulations and highlights the importance of the genetic architecture of climate adaptation.

### Introgression and extinction

Here we’ve focused on the possibility of species collapse, but another more common predicted effect of climate change is extinction. In our model, population sizes are constant and fitness is relative, so extinction is impossible. Despite this, our results do have implications for the likelihood of extinction.

If RI alleles contribute to local adaptation (i.e. are extrinsic), then the collapse of RI must result in the spread of fitness reducing maladaptive non-local loci introgress, similar to the concept of linkage drag in plant breeding (Zamir 2001). The net effect of the introgression is positive in our climate change scenario, since the maladaptive introgressed alleles are linked to positive climate QTL alleles, but the overall fitness of the population is lower than it would be if it adapted *without* introgression. In this way, adaptive introgression makes populations better adapted to the changing climate, but less adapted to their home niche. In the real world, this may be reflected in reduced population size or growth rate and could increase the chance of extinction. This effect is dependent on the amount and type of RI loci swept to fixation by linkage with beneficial climate QTL, which in itself will depend on the speed of fixation and amount of recombination. Thus stronger selection, smaller population size or reduced recombination rate will all increase linkage drag. Note that if RI is intrinsic, this doesn’t necessarily apply because intrinsic RI loci are only detrimental based on interactions with other loci and won’t have any fitness penalty if all RI alleles are homogenized by introgression.

On the other hand, it is likely that the adaptive introgression of climate QTL increases the chances that a species can adapt to a changing environment instead of going extinct. Again, we can’t directly address this question in our model but we do see that when all introgression is prevented adaptational lag increases (Figure 4). This is consistent with the larger total gene pool of adaptive variants available when gene flow is possible. The magnitude of this effect will depend on the amount and architecture of divergently selected loci (i.e. the potential linkage drag) as well as the diversity of climate QTL alleles in each species (i.e. the effective population size).

### Linkage and the genetic architecture of climate adaptation

A key aspect of our model is that while RI loci occur at predefined intervals in the genome, climate-sensitive alleles can arise at any other locus in the genome. This allows for climate-sensitive alleles to become readily linked to RI-causing alleles and eventually introgress if the combined effect of positively selected climate alleles exceeds the deleterious effect of the linked RI allele. The incidental establishment of this linkage within the two adapting populations is a fundamental cause of later introgressive collapse. This is supported by our simulations that varied the number of RI loci and also incidentally varied the average degree of linkage between all climate-sensitive loci and all RI loci. We found that in simulations with more RI loci, and therefore a higher probability of linkage between RI and climate-sensitive loci, there was greater loss of reproductive isolation. Thus, a key question is whether such linkage could plausibly be established in a natural population.

Several lines of evidence suggest that this is likely to be true. First, the genetic architecture of adaptation to a changing climate is likely to closely resemble the architecture of local adaptation in general, i.e. a large number of small effect alleles with a smaller number of large effect loci (reviewed in Savolainen et al. 2013). This idea is directly supported by recent work showing that climatic adaptation in conifers is underlain by large number of loci scattered throughout the genome, with the majority of these showing modest phenotype-environment correlations (Yeaman et al. 2016). Secondly, recent analyses of large human datasets support the idea that most complex traits (of any kind) are probably determined by a large number of small-effect loci found nearly everywhere in genome along with a handful of “core genes” (Boyle et al. 2018). Thus, given that the genetic architecture of RI is itself likely to be highly polygenic (further discussed in Ravinet et al. 2017), it seems highly plausible that linkage between climate-sensitive alleles and RI alleles can readily occur in natural populations.

### The role of recombination rate

Recombination rate is thought to play a key role in mediating patterns of divergence and introgression in natural populations (e.g. Samuk et al. 2017, Schumer et al. 2018). Specifically, regions of high recombination are thought to be less resistant to gene flow because of decreased linkage between alleles conferring RI. We see this in our simulations, as simulations with higher recombination rates have greater amounts of introgression in both control and climate change scenarios. Interestingly, we observe a slight negative correlation between recombination rate and loss of RI under the climate change scenario (Figure 3, Figure S1). This effect is a direct consequence of lower recombination rates leading to increased linkage between RI alleles and globally-adaptive climate alleles. This increased linkage leads in turn to larger numbers of RI alleles being dragged along with globally-adaptive alleles and homogenized between populations. While mutations were not particularly limiting in our simulations, adaptive introgression in regions of low recombination should generally require larger effect alleles than introgression in regions of high recombination, particularly at the onset of climate change when selection is weaker overall.

### The strength of climate-mediated selection

A strong shared selection pressure is ultimately the key mediator of the collapse of RI we observed. Was the magnitude of simulated selection necessary to cause this collapse realistic? One way to assess this is to measure the magnitude of the phenotypic response to selection in our simulations and compare it to estimates from natural systems. In our case, the phenotypic response to selection ranged from 0.01-0.06 Haldanes (standard deviations per generation). This is in line with the magnitude of phenotypic response observed in both natural and anthropogenically-induced selection (e.g. Hendry et al. 2008). Further, this is below the theoretical threshold of 0.1 Haldanes thought to result in an unsustainable long-term response to selection (for N_e_ = 500; Lynch & Lande, 1993; Bürger & Lynch, 1995).

Another way of assessing the realism of our scenarios is comparing the selection coefficients of climate QTL in our simulations with values measured in empirical studies. We measured selection coefficients in our example simulation by comparing relative fitness values for samples with and without each mutation (Appendix 1), and therefore are capturing not just the effect of the climate QTL mutation, but also all linked loci. Only 0.7% of introgressed climate QTL loci had selection coefficients > 1, within the range of empirical values, therefore adaptive introgression is occurring without abnormally high selection coefficients (Kingsolver et al. 2001).

Thus, the strength of selection we modelled was not particularly extreme nor would it necessarily lead to the extinction of the modelled populations. It is also worth noting that the estimated rate of phenotypic change in wild populations due to future GCC is thought to be at least as large as the rates we described here, and are projected to likely exceed 0.1 Haldanes in many cases (Gienapp, Leimu, & Merilä, 2007; Merilä & Hoffman 2016). In sum, the global strength of phenotypic selection simulated here was not unrealistically high, and if anything represents a conservative adaptive scenario.

## Conclusion

Hybridization is a double-edged sword under rapid environmental change. It can allow species access to a larger pool of adaptive alleles but linkage with RI alleles will weaken overall RI and may lead to speciation reversal. Importantly, our work highlights the dangers of hybridization for a much wider pool of species, not just those on range margins or with existing porous species boundaries. In the longer term, we predict that specific cases of speciation reversal should be linked to climate change but we also predict effects before full speciation reversal. If our model is correct, we predict that alleles conferring adaptation to present and future climate (e.g. heat or drought) will be more likely to introgress between species. Although identifying all the loci underlying climate adaptation is challenging, recent work by Exposito-Alonso et al. (2019) highlights that it can be done. Such an approach can be combined with sequencing data in related species to identify where introgression is occuring. Our results also suggest that hybrid zones should become increasingly porous as climate adaptation alleles move between species and that this effect would be stronger in regions with more dramatic climate change (e.g. the arctic). This could be done by resampling previously studied hybrid zones or by comparing contemporary samples to museum and herbarium samples. Confirmation of these predictions would show that climate adaptation is occurring through a larger multi-species gene pool and be a warning sign for the future homogenization of these species.

## Acknowledgements

This work was supported by an NSERC Banting Postdoctoral Fellowship to GLO and an NSERC Postdoctoral Fellowship to KS. KS was further supported by postdoctoral funding and good vibes from Mohamed Noor at Duke University. Matthew Osmond, Kate Ostevik and Loren Rieseberg provided helpful feedback and discussions on earlier versions of the manuscript. We thank Philip Messer and Ben Haller for assistance with the SLiM 3.X software.

## Supplemental Figures

**Figure S1.**
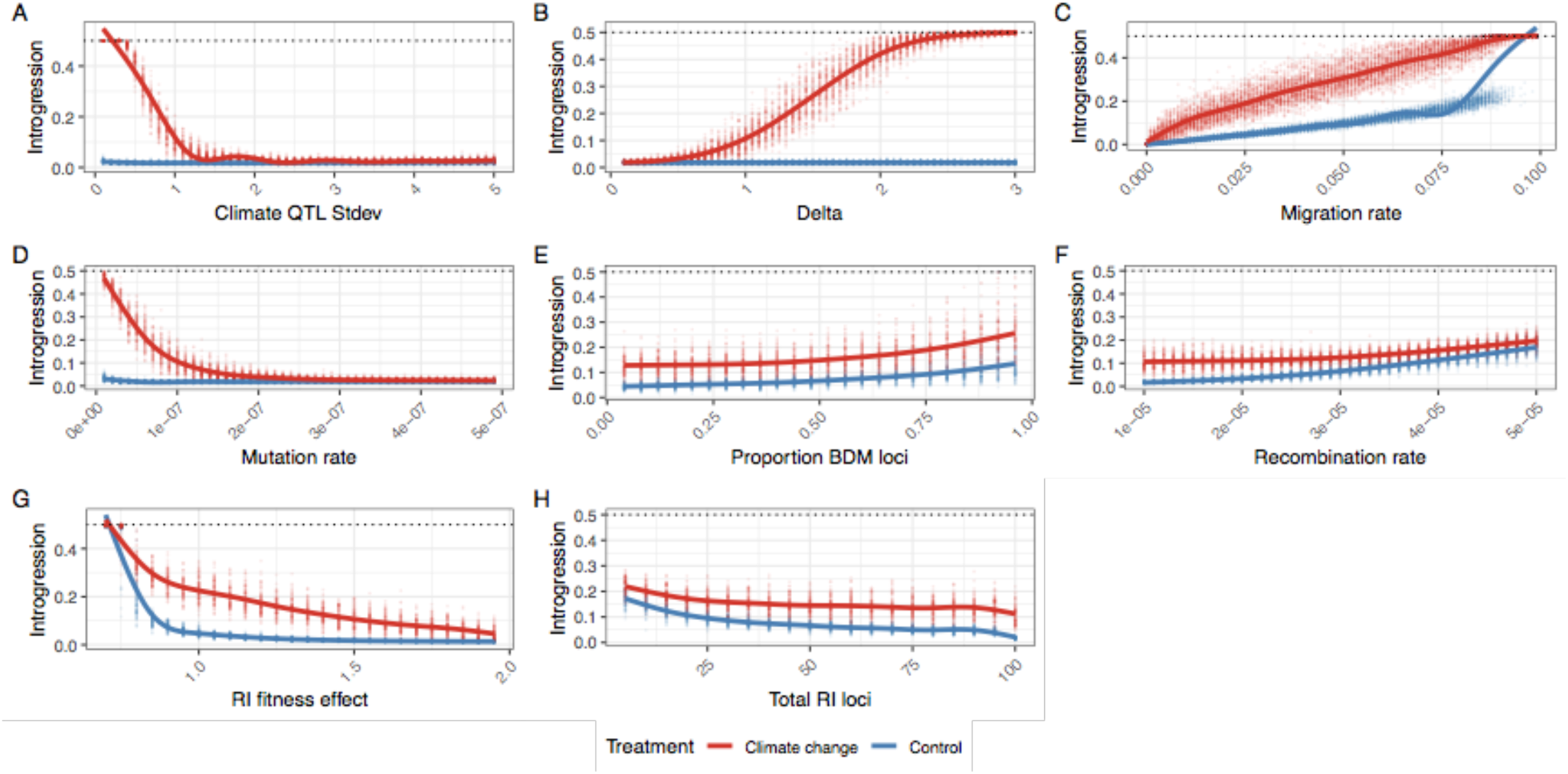
The average amount of introgression at generation 10,100 for climate change (red) and control simulations (blue), while varying individual parameters. Complete homogenization of both populations occurs when average introgression = 0.5. Individual parameters were varied to show the effect of (A) climate QTL effect size standard deviation, (B) optimum shift per generation (delta), (C) migration rate, (D) climate QTL mutation rate, (E) proportion of RI loci that are BDM instead of extrinsic, (F) the recombination rate, (G) the fitness effect of each RI loci and (H) the number of RI loci.

**Figure S2.**
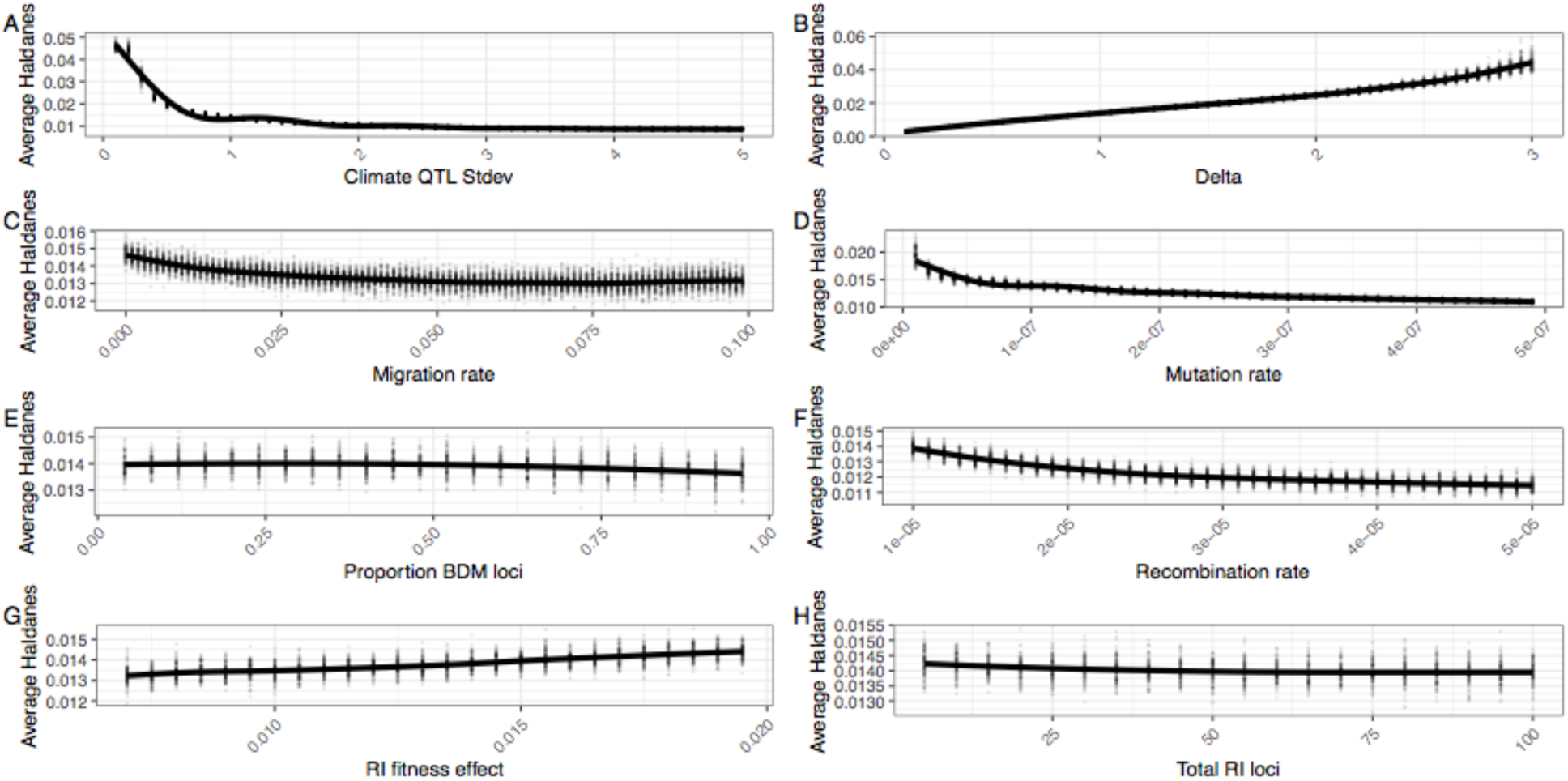
The average Haldanes for the post-burn in period with climate change. Individual parameters were varied to show the effect of A) climate QTL effect size standard deviation, (B) optimum shift per generation (delta), (C) migration rate, (D) climate QTL mutation rate, (E) proportion of RI loci that are BDM instead of extrinsic, (F) the recombination rate, (G) the fitness effect of each RI loci and (H) the number of RI loci. Note that Haldanes are scaled by trait variation, so the same rate of absolute phenotypic change (i.e. Darwins) can have multiple different Haldanes if the phenotypic variability changes. In all panels except B, the rate of phenotypic change roughly matches the rate of environmental change, so the differences in Haldanes reflects differences in trait variance. Lower trait variance (e.g. through reduced mutation rate or lower QTL standard deviation) produces higher haldanes for the same absolute rate of phenotypic change.

