## Supplementary Appendix 1 for "Adaptive introgression during environmental change can weaken reproductive isolation"

### **Appendix 1:** Equations used in simulations and analyses

*Fitness calculations*

Fitness in this model is determined by how well the sample's phenotype matches the current climate optimum as well as the genotypes of reproductive isolation alleles. During the initial burn-in, the climate optimum oscillates slightly above and below zero with an amplitude of $a$ and a frequency of $f$ Eq. 1, and then increases linearly during the climate shift period by $\Delta$ units per generation Eq. 2.

$$(1)\ O_{g_{burn}} = sin(\pi * gen/f)a$$
$$(2)\ O_{g_{shift}} = \Delta(g - b)$$

The fitness effect of the climate match is calculated by first calculating each sample's climate phenotype by summing the effect sizes, Q, of each copy of a QTL allele present in the sample. In this way, QTLs are all additive and co-dominant (dominance=0.5). The fitness effect of this phenotype is determined by a gaussian function with a mean of the current climate optimum $O_g$ and a standard deviation of $sd_{climate}$ Eq. 3. Samples with climate phenotypes distant from the optimum have reduced relative fitness.

$$(3)\ \omega_{climate} = f\left(\left(\sum_{i=1}^{n_{sample\ QTL}} Q_i\right) - O_g \mid 0,\ sd_{climate}\right)$$

Along with climate adaptation, fitness is also determined by the alleles at reproductive isolation loci which are either extrinsic or intrinsic epistatic. In most simulations, all RI loci are extrinsic, except in simulations designed to test the effect of Bateson-Dobzhansky-Muller (BDM) incompatibilities. The effect of extrinsic RI loci is determined by summing the counts of non-local alleles divided by 2 (for co-dominance) Eq. 4.

$$(4)\ n_{EX} = \prod_{1}^{l_{EX}} \frac{g_{AWAY}}{2}$$

BDM loci were initialized as randomly selected pairs. In each pair, both populations are initially fixed for different alleles in the simulation. One locus is set as the derived state ($A$) in population 1 and the other as the derived state ($B$) in population 2. Negative interactions occur when both derived alleles are present in a single diploid individual and are equally deleterious in all combinations (Table S1). Thus each BDM pair can produce epistasis $(\tau)$ counted as 0 or 2, and this is summed for each individual Eq. 5.

**Table S1** | Genotype epistasis $(\tau)$ for BDM loci

| | *aa* | *Aa* | *AA* |
|---|---|---|---|
| *bb* | 0 | 0 | 0 |
| *Bb* | 0 | 2 | 2 |
| *BB* | 0 | 2 | 2 |

$$(5)\ n_{BDM} = \prod_{i=1}^{l_{BDM}/2} \tau_i$$

The total sum of divergently selected alleles $n_{EX}$ and BDM epistasis $n_{BDM}$ are treated as independent alleles with selection coefficients of $s_{RI}$ and are multiplicatively added Eq. 6. Importantly, this puts BDM and extrinsic alleles on the same scale, so they are comparable. Although this model results in diminishing returns in terms of absolute fitness, relative fitness scales correctly.

$$(6)\ \omega_{RI} = (1 - s_{RI})^{(n_{EX}+n_{BDM})}$$

The two measures of fitness are combined to create the fitness measure of each sample Eq. 7.

$$(7)\quad \omega = \omega_{climate} * \omega_{RI}$$

*Reproductive isolation calculations*

To see if rapid shifts in the phenotypic optimum can lead to reverse speciation, we measured average reproductive isolation during the climate shift period. We operationally defined reproductive isolation as the difference in fitness between an average migrant individual vs. an average non-migrant (i.e. the difference in expected "home" vs. "away" fitness). This is determined purely based on the extrinsic and BDM RI alleles, and does not include climate QTL alleles.

For extrinsic loci, we can think of the reproductive isolation in terms of a representative individual being transported from its own population to the other population and measuring its relative fitness. We calculate average "home" fitness by measuring the proportion of foreign alleles and applying it to Eq. 6 to get an expected fitness penalty. This is averaged for both populations Eq. 8.

$$(8)\ \ E\overline{\varpi}_{EX_{HOME}} = \frac{(1-s_{RI})^{(p_{EXp1_{AWAY}} * l_{EX})} + (1-s_{RI})^{(p_{EXp2_{AWAY}} * l_{EX})}}{2}$$

“Away” fitness was calculated in a similar way, but using the proportion of home alleles when calculating the fitness penalty Eq. 9.

$$(9)\ \ E\overline{\varpi}_{EX_{AWAY}} = \frac{(1-s_{RI})^{(p_{EXp1_{HOME}} * l_{EX})} + (1-s_{RI})^{(p_{EXp2_{HOME}} * l_{EX})}}{2}$$

Reproductive isolation from BDMs are more complicated because BDMs have no effect in generation 0 after migration (i.e. before breeding). Their effect only appears after mating in F1s and beyond which have combinations of alleles from both populations. Thus, for BDMs, we estimate expected RI based on Hardy-Weinberg expectations of genotype frequencies assuming mating within the population, and mating between populations (Tables S2 & S3). From this expectation, we estimated the expected amount of BDM epistasis (Eq 10 & 11). These formulae assume that BDM loci interacting pairs are unlinked and segregate independently.

**Table S2** | Expected genotype frequencies for interpopulation crosses.

| | $aa$ | $Aa$ | $AA$ |
|---|---|---|---|
| $bb$ | $a_1a_2b_1b_2$ | $(A_1a_2 + a_1A_2)b_1b_2$ | $A_1A_2b_1b_2$ |
| $Bb$ | $a_1a_2(B_1b_2 + b_1B_2)$ | $(A_1a_2 + a_1A_2)(B_1b_2 + b_1B_2)$ | $A_1A_2(B_1b_2 + b_1B_2)$ |
| $BB$ | $a_1a_2B_1B_2$ | $(A_1a_2 + a_1A_2)B_1B_2$ | $A_1A_2B_1B_2$ |

**Table S3** | Expected genotype frequencies for intrapopulation crosses.

| | $aa$ | $Aa$ | $AA$ |
|---|---|---|---|
| $bb$ | $a^2b^2$ | $2(Aa)(b^2)$ | $A^2b^2$ |
| $Bb$ | $2(a^2)(Bb)$ | $4(Aa)(Bb)$ | $2(A^2)(Bb)$ |
| $BB$ | $a^2B^2$ | $2(Aa)(B^2)$ | $A^2B^2$ |

$$(10)\ E\varpi_{BDM_{HOME}} = \frac{\sum_{p1}^{p2} \sum_{i=1}^{n_{BDM\ pairs}} (2*E(AABB_i))+(2*E(AABb_i)) + (2*E(AaBB_i))+(2*E(AaBb_i))}{2}$$

$$(11)\ E\varpi_{BDM_{AWAY}} = \sum_{i=1}^{n_{BDM\ pairs}} (2*E(AABB_i)) + (2*E(AAbB_i)) + (2*E(AaBB_i)) + (2*E(AaBb_i))$$

After all average fitness values are compiled, an estimated RI score is calculated using Eq. 12. This RI score represents the *average fold higher fitness in the home population compared to the other (away) population*. If there is no RI or populations are completely admixed, RI will equal 1, representing equal fitness in either environment.

$$(12)\ \overline{RI} = (1 - s_{RI})^{(E\varpi_{BDM_{AWAY}} + E\varpi_{EX_{AWAY}})} / (1 - s_{RI})^{(E\varpi_{BDM_{HOME}} + E\varpi_{EX_{HOME}})}$$

*Selection coefficient visualization*

To understand the scale of fitness effects in the simulation, we collected individual genotype values and sample fitness for each sample in each generation during the climate shift period (i.e. generation 10001 to 10100). We then calculated $s$ using Equation 13. This equation compares the mean fitness for samples homozygous for the mutation with samples homozygous for the wild-type allele and is normalized by the mean fitness for wild-type samples. We required that each group needed to have at least 2 individuals for $s$ to be calculated.

$$(13)\ s = (\overline{w}_{11} - \overline{w}_{00})/\overline{w}_{00}$$

In this case, $s$ is the realised fitness effect which includes the effect of linked loci. We plotted mutations with intermediate allele frequencies (0.1 < frequency < 0.9) because at frequencies closer to 0 and 1, mutations are most often found in first generation migrants, and have more extreme variation in fitness (Figure 2D).

**Table S4** | List of parameters used in formulas.

| Symbol | Parameter |
|---|---|
| *gen* | Generation number |
| *b* | Burn in generations |
| *f* | Burn in oscillation wavelength |
| *a* | Burn in oscillation amplitude |
| Q | QTL loci strength |

| | |
|---|---|
| $O$ | Climate optimum |
| $sd_{climate}$ | Standard deviation of gaussian climate fitness function |
| $s_{RI}$ | Selection at individual divergently selected or BDM loci |
| $s$ | Realized selection coefficient |
| $l$ | Total number of loci |
| $n$ | Number of alleles in sample |
| $\Delta$ | Change in climate optimum per generation during shift |
| $g$ | Sample genotype |
| $\omega$ | Fitness effect |
| $\overline{w}$ | Average realized fitness |
| $\tau$ | Epistasis effect |
| $E\,\overline{\omega}$ | Average estimated fitness score |
| $\overline{RI}$ | Reproductive isolation score |
| $N$ | Population size |
| $p$ | Allele frequency |
